# Quantification of elongation stalls and impact on gene expression in yeast

**DOI:** 10.1101/2023.03.19.533377

**Authors:** Wanfu Hou, Vince Harjono, Alex T. Harvey, Arvind Rasi Subramaniam, Brian M. Zid

**Affiliations:** Department of Chemistry and Biochemistry, University of California San Diego, La Jolla, CA, USA; Basic Sciences Division and Computational Biology Section of Public Health Sciences Division, Fred Hutchinson Cancer Center, Seattle, WA, USA

**Keywords:** translation elongation, ribosome stalling, codon optimality, ribosome quality control, no-go decay

## Abstract

Ribosomal pauses are a critical part of co-translational events including protein folding and localization. However, extended ribosome pauses can lead to ribosome collisions, resulting in the activation of ribosome rescue pathways and turnover of protein and mRNA. While this relationship has been known, the specific threshold between permissible pausing versus activation of rescue pathways has not been quantified. We have taken a method used to measure elongation time and adapted it for use in *S. cerevisiae* to quantify the impact of elongation stalls. We find, in transcripts containing Arg CGA codon repeat-induced stalls, a Hel2-mediated dose-dependent decrease in protein expression and mRNA level and an elongation delay on the order of minutes. In transcripts that contain synonymous substitutions to non-optimal Leu codons, there is a decrease in protein and mRNA levels, as well as similar elongation delay, but this occurs through a non-Hel2-mediated mechanism. Finally, we find that Dhh1 selectively increases protein expression, mRNA level, and elongation rate. This indicates that distinct poorly translated codons in an mRNA will activate different rescue pathways despite similar elongation stall durations. Taken together, these results provide new quantitative mechanistic insight into the surveillance of translation and the roles of Hel2 and Dhh1 in mediating ribosome pausing events.

---

Production of cellular proteins through translation is crucial for maintaining homeostasis and adapting to changing environmental conditions. Translation can be broken down into three sequential steps: initiation, during which ribosomes assemble at the initiation site on an mRNA, elongation, during which ribosomes translocate across the mRNA and build upon a nascent peptide, and termination, during which ribosomes are removed from the mRNA, recycled, and the newly synthesized protein is released. Cells dedicate many resources to the monitoring, regulation, and quality control of protein synthesis as dysregulation may lead to aberrant cellular function and neurological diseases such as ALS (1, 2).

Each step in the translation process is governed by various regulatory steps. Initiation has long been known to be the rate-limiting step in translation for most mRNAs and subject to intense regulation (3, 4). Recent studies have focused on elongation as another important regulatory step in protein synthesis (5). Indeed, modulation of elongation speed has been shown to serve a functional role in both proper protein folding (6–10) and localization (11–14). These examples give credence to the notion that ribosome pausing is essential for certain cellular processes. Recent reports using ribosome profiling to analyze disome peaks have estimated that upwards of 10% of translating ribosomes are engaged in the disome state (9, 15, 16), indicating the commonplace occurrence of ribosome collisions.

The functional and necessary nature of ribosome stalls, however, makes it challenging for cellular machinery to distinguish between beneficial stalls and situations requiring ribosome rescue. Ribosomes that undergo translation on an aberrant mRNA, such as on a truncated mRNA, stall in place and are unable to be disassembled by translation termination machinery (17–19). Upon extended stalling events, translating ribosomes may collide with stalled ribosomes, resulting in a ribosome collision which can eventually lead to further accumulation of collided ribosomes. Two pathways may be activated upon the detection of these collision events: (1) ribosome quality control (RQC), which leads to the rescue and recycling of stalled ribosomes, and (2) no-go decay (NGD), which leads to the endonucleolytic cleavage and subsequent degradation of the aberrant transcript (20–22). Both pathways are triggered by the ribosome collision sensor Hel2(yeast)/ZNF598(mammals) which detects disome formations which form as a result of prolonged ribosome stalling (23). In RQC, Hel2 ubiquitinates the small ribosomal subunit which leads to activation of the RQC trigger (RQT) complex in yeast, ultimately resulting in ribosome disassembly and degradation of the nascent peptide (17–19). Concurrently, Hel2 activation also leads to the activation of the NGD pathway which results in mRNA degradation primarily through the endonuclease Cue2 and the exonucleases Xrn1 and Ski7 (19, 24, 25). Hel2 and other sensors of elongation quality must maintain a balance between permitting transient and functional stalls while at the same time engaging rescue pathways to prevent the buildup of ribosomes on problematic mRNA.

How elongation quality sensors can distinguish between functional stalls and those requiring rescue pathways remains unclear. It has been proposed that the severity of ribosome collision may determine which cellular response is activated in response to a collision event (26). Supporting this model, a recent study by Goldman and colleagues found that clearance of stalled ribosomes was far slower than elongation and termination and proposed that slow ribosome clearance allows cells to distinguish between transient and deleterious stalls (27). While this model may explain how functional stalls and detrimental stalls either resume elongation or initiate RQC using the same surveillance pathways, respectively, the definition of “severity” in this context remains vague. Are the distinguishing factors the time duration of the stall, the number of ribosome collisions (27), the specific location and context of where the stall occurs on a transcript, or a combination of all these factors and more? It is from this lack of understanding of how cellular surveillance machinery can distinguish between these two opposing outcomes that necessitates reliable, quantitative methods to describe the various aspects of ribosome stalling events.

One important factor that contributes to elongation speed is codon optimality, a metric that describes the translational efficiency of the 61 amino acid specifying codons. Codon optimality, unique to each species, takes into account various factors implicated in elongation rate, including tRNA availability and demand, frequency of use in the genome, GC content, and interactions with the ribosome exit tunnel (28– 31). Furthermore, codon optimality has been found to correlate with elongation speed and mRNA decay, with transcripts enriched in “optimal” codons associated with faster elongation speed and lower mRNA decay rates and those enriched in “nonoptimal” codons associated with slower elongation speed and higher mRNA decay rates (30, 32–41). While many studies, both in vivo and in vitro, have assessed the impact of synonymous codon substitutions on protein expression, mRNA decay, and ribosome pausing, quantification of the impact on elongation time has not been widely available.

In this study, we describe the development of an *in vivo* quantitative luciferase-based assay to measure elongation time. We assessed the time delay associated with acute stalls caused by the inclusion of repeats of the nonoptimal arginine codon CGA and find that elongation time increases in a dose-dependent manner. Surprisingly, we find that no go RNA decay reaches a maximum level at a specific stall length despite increasing translation elongation times and protein expression continuing to decrease. Furthermore, we assessed the effect of synonymous codon substitutions on elongation time of a standardized ORF and identified the non-optimal leucine codon CTT as a strong driver of elongation delay. CTT’s effect on elongation time where context dependent as only specifically localized non-optimal CTT’s caused significant elongation delays and protein repression. The development of this assay and our findings provide steps towards a detailed understanding of the triggers of ribosome quality control pathways.

## Results

### Development and validation of elongation assay

To create a quantitative elongation duration reporter assay, we utilized a tetracycline-inducible promoter to control mRNA induction of a bioluminescent nanoluciferase (nLuc) reporter downstream of open reading frames (ORFs) of interest. The nLuc reporter has been previously studied in yeast under the control of a stress-inducible promoter and its bioluminescent output faithfully recapitulates induced mRNA levels after heat shock (42). To test this system, we developed a series of constructs in which we varied the length of the upstream ORF by insertion of yeast-optimized yellow fluorescent protein (YFP) or yeast-optimized monomeric infrared red fluorescent protein (miRFP) ORFs upstream of nLuc (Figure 1A). Tet-nLuc is included to control for the time cost of initiation steps including anhydrotetracycline (ATC) penetration, transcription initiation, mRNA export, and translation initiation. nLuc protein expression was collected for each construct over 60 minutes and normalized to OD600 measured at T=0 min (time of ATC addition). Elongation time was calculated using a Schleif plot (43) and adjusted based on an average mRNA transcription time of 1500 nucleotides per minute (14, 44). We find a delay in the first appearance of nLuc upon the addition of *optYFP* and a further delay in the longer *miRFP-optYFP-nLuc* reporter (Figure 1B). We then used these measured delays to calculate the translation elongation rate of *optYFP* and *miRFP* ORFs as approximately 4 AA/sec and 3 AA/sec (Figure 1C), respectively, which is consistent with bulk elongation rate measurements of 3-10 AA/sec (45, 46). We do not find a significant difference in elongation rate between the two optimized ORFs. This implies that our reporter can quantify the in vivo translation rates of our reporters.

**Fig. 1.**
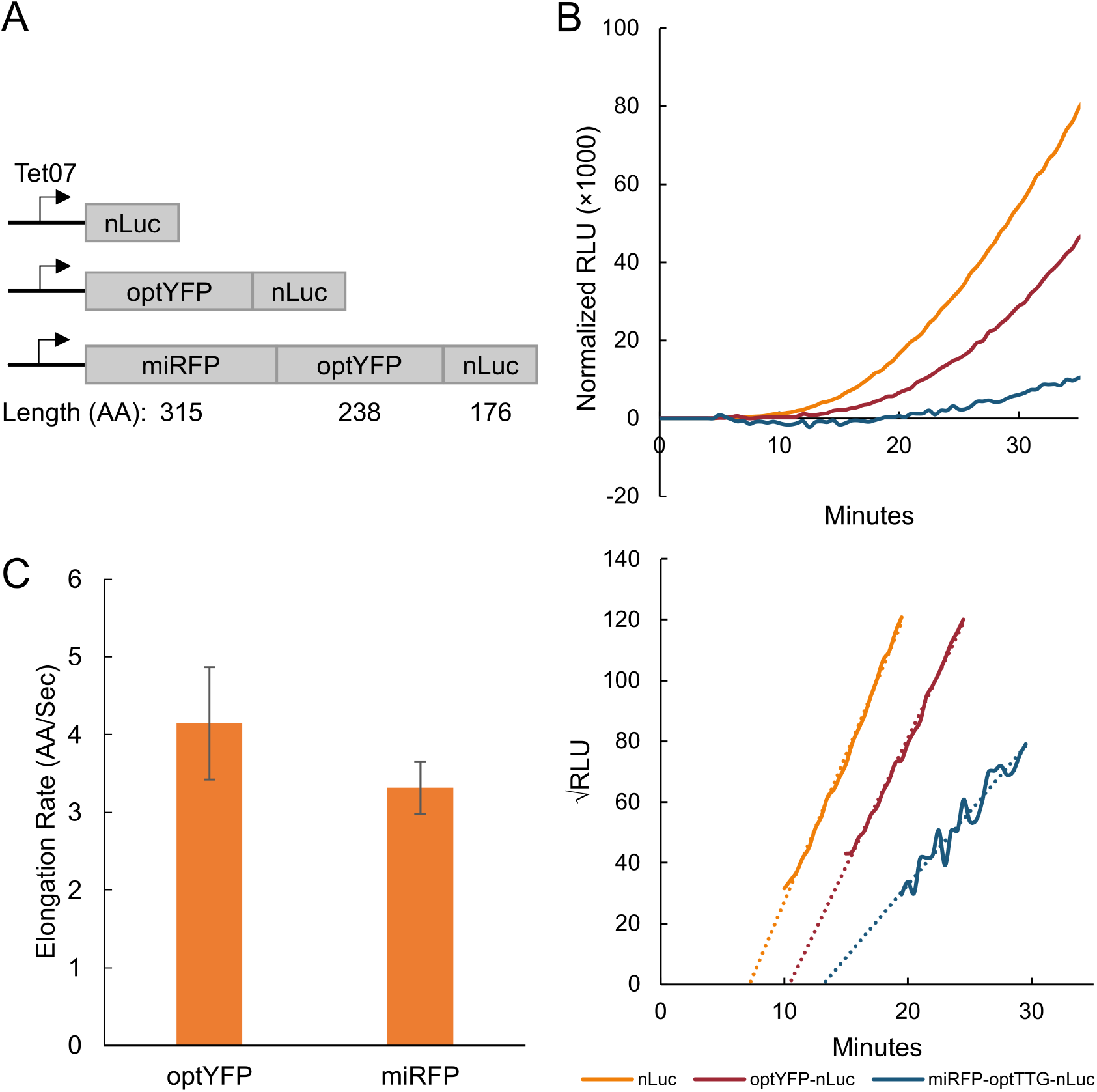
Assay validation via elongation rate measurements. A: Diagram of yeast-optimized constructs of various lengths. Optimized YFP (optYFP) or both optYFP and optimized miRFP (miRFP) are set upstream of a nanoluciferase (nLuc) reporter. Constructs are expressed from an inducible Tet07 promoter. B: (Top) Representative assay data of relative light units (RLU) of each construct over time normalized to OD600. (Bottom) Schleif plot and associated trendlines of the top graph. C: Calculated elongation rate measurements of optYFP (n=9) and miRFP (n=4) ORFs. Error bars indicate SEM.

### Hel2 decreases protein expression, mRNA levels, and delays elongation in acute CGA constructs

To further explore the utility of our reporter we wanted to verify this system could quantify the duration of elongation pauses of known ribosome stalling sequences. Consecutive non-optimal CGA arginine codons are known to induce slow translation elongation and terminal stalling through wobble decoding of CGA (47–50). To quantify the effect of these non-optimal codons on elongation time and gene expression, we developed a series of constructs in which we inserted between 2 and 6 tandem CGA repeats between the yeast-optimized YFP ORF and nLuc reporter ORF shown previously (Figure 2A). First, we tested the protein expression of our induced constructs and found a dose-dependent exponential decline in protein production as the number of CGA codons increased, similar to a previous study by Letzring and colleagues (48) (Figure 2B). We however, did not see a significant impact on protein expression until 3 CGA codons were included. Next, we measured mRNA levels and found that mRNA levels significantly decreased with the addition of 3 CGA codons but mRNA levels remained constant around 40% of our control construct regardless of additional CGA codons (Figure 2C). We then measured the elongation delay in each of our constructs by comparing to a control reporter lacking any CGA codons (Figure 2D). We found that elongation delay increased in a dose-dependent manner beginning at 3xCGAs, with 6xCGA causing an 4.5 minute extension of the translation duration. There was a relatively linear relationship between CGA stall number after 3 CGAs and elongation time which allowed us to calculate that each CGA adds approximately 76 seconds to the overall elongation time. We found that this elongation delay was specifically due to CGA codons as a 6x AGA codon, which also encodes for arginine, had no effect on elongation time (Supplementary Figure 1).

**Fig. 2.**
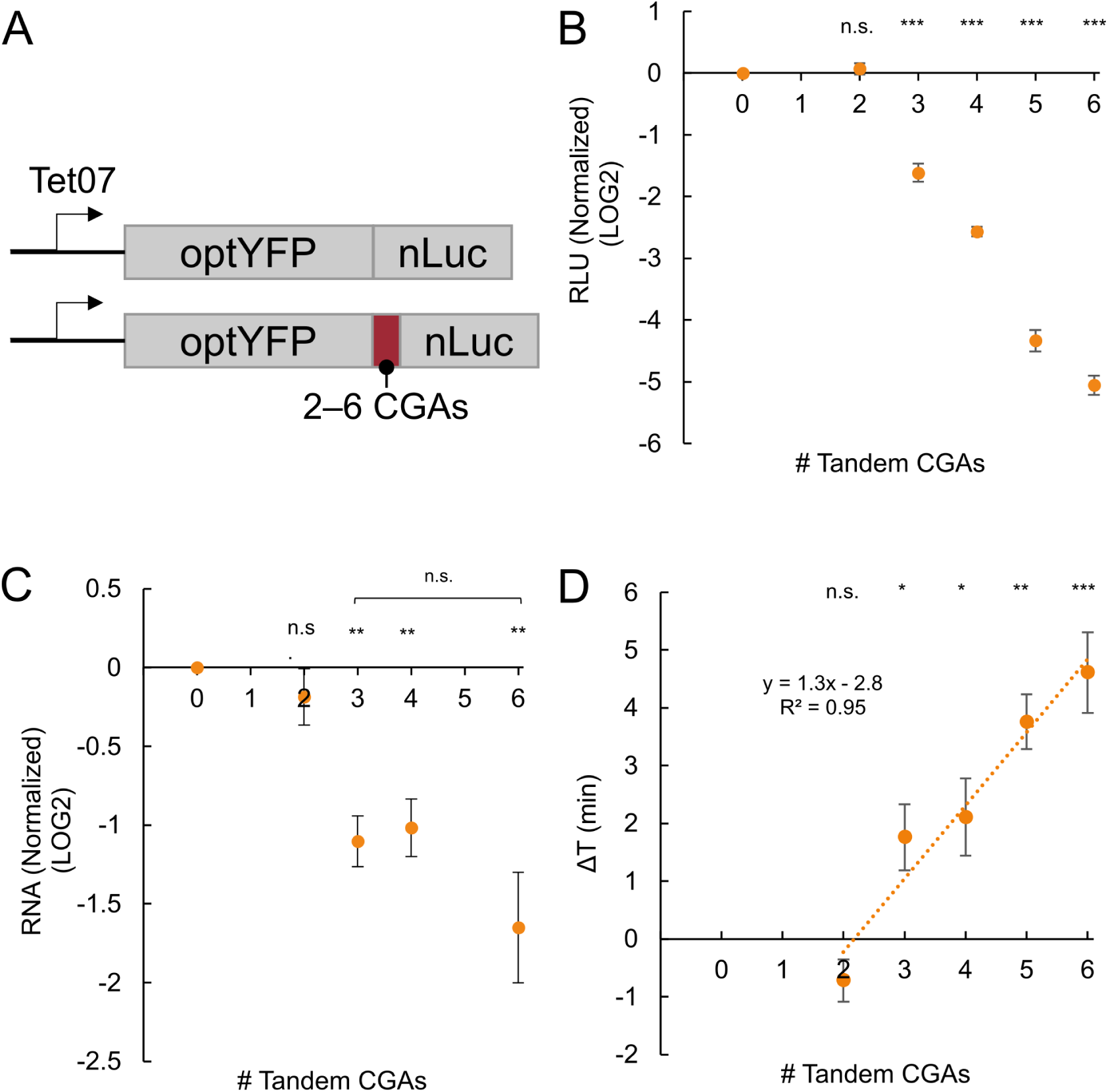
CGA-derived acute stalls negatively impact gene expression and increase elongation time in a dose-dependent manner. A: Diagram of optimal and CGA-containing constructs. Between 2 and 6 CGAs are inserted between the optYFP and nLuc ORFs. B: Protein expression of CGA constructs at T=60 min normalized to optimized control (2xCGA n=10, 3xCGA n=8, 4xCGA n=10, 5xCGA n=5, 6xCGA n=10). C: mRNA levels of CGA constructs at T=60 min normalized to optimized control. (n=3). C: Elongation delay of CGA-containing constructs compared to optimized control. (n=3). All error bars indicate SEM. All statistical significances were calculated for each construct using two-tailed paired Student’s t-Test against optYFP control.

We then tested the role of Hel2 and Syh1, two factors implicated in impacting gene expression due to prolonged ribosome stalls. Hel2 is a translation surveillance factor that senses ribosome collisions and activates the ribosome rescue pathways ribosome quality control (RQC) and no-go decay (NGD) pathways which result in protein and mRNA turnover, respectively. Syh1 is a homolog of the mammalian NGD factor GIGYF1/2 that was previously found to have a role in NGD in yeast (50, 51). We measured protein expression in our constructs containing 2, 4, and 6 CGAs in a *hel2*Δ and *syh1*Δ background and we compared it to their wild type (WT) counterparts (Figure 3A, Supplementary Figure 2A). We found that deletion of *HEL2* partially rescued protein expression in the 4xCGA and 6xCGA, but *SYH1* deletion had no effect on the 6xCGA protein expression. We next measured RNA levels in our 2xCGA, 4xCGA, and 6xCGA strains and found that RNA levels were increased in our 4xCGA and 6xCGA-containing *hel2*Δ strains but there was no change in the 2xCGA strain (Figure 3B). We found no significant difference in RNA levels for the 6xCGA in our *syh1*Δ strain (Supplementary Figure 2B). Together, these results imply Hel2-mediated RQC and NGD are partially responsible for the observed decrease in protein and RNA levels, respectively, in the wild-type strains. Lastly, we sought to measure the impact of Hel2 on elongation time. A recent review by Meydan and Guydosh proposed two non-mutually exclusive models of Hel2’s activity on the stability of ribosome collisions: (1) Hel2 is necessary to rescue stalled ribosomes and *hel2* deletion would result in further buildup of collided ribosomes and (2) Hel2 stabilizes collided ribosomes and *HEL2* deletion would result in reduced ribosomal pausing (26). Model 2 that Hel2 stabilizes collided ribosomes was further supported by experimental data that *hel2*Δ reduces disome pauses in ribosome profiling data sets (52). To assess the effect of Hel2 on elongation time and distinguish between these two models, we compared the elongation time of our control, 4xCGA, and 6xCGA strains between WT and *hel2*Δ backgrounds and found no significant difference in our control strain but a decrease in overall elongation time in our 4xCGA and 6xCGA strains when expressed in a *hel2*Δ background (Figure 3C). This suggests that Hel2 functions to slow down elongation in our CGA-containing strains and is consistent with the second proposed model in which Hel2 stabilizes collided ribosomes.

**Fig. 3.**
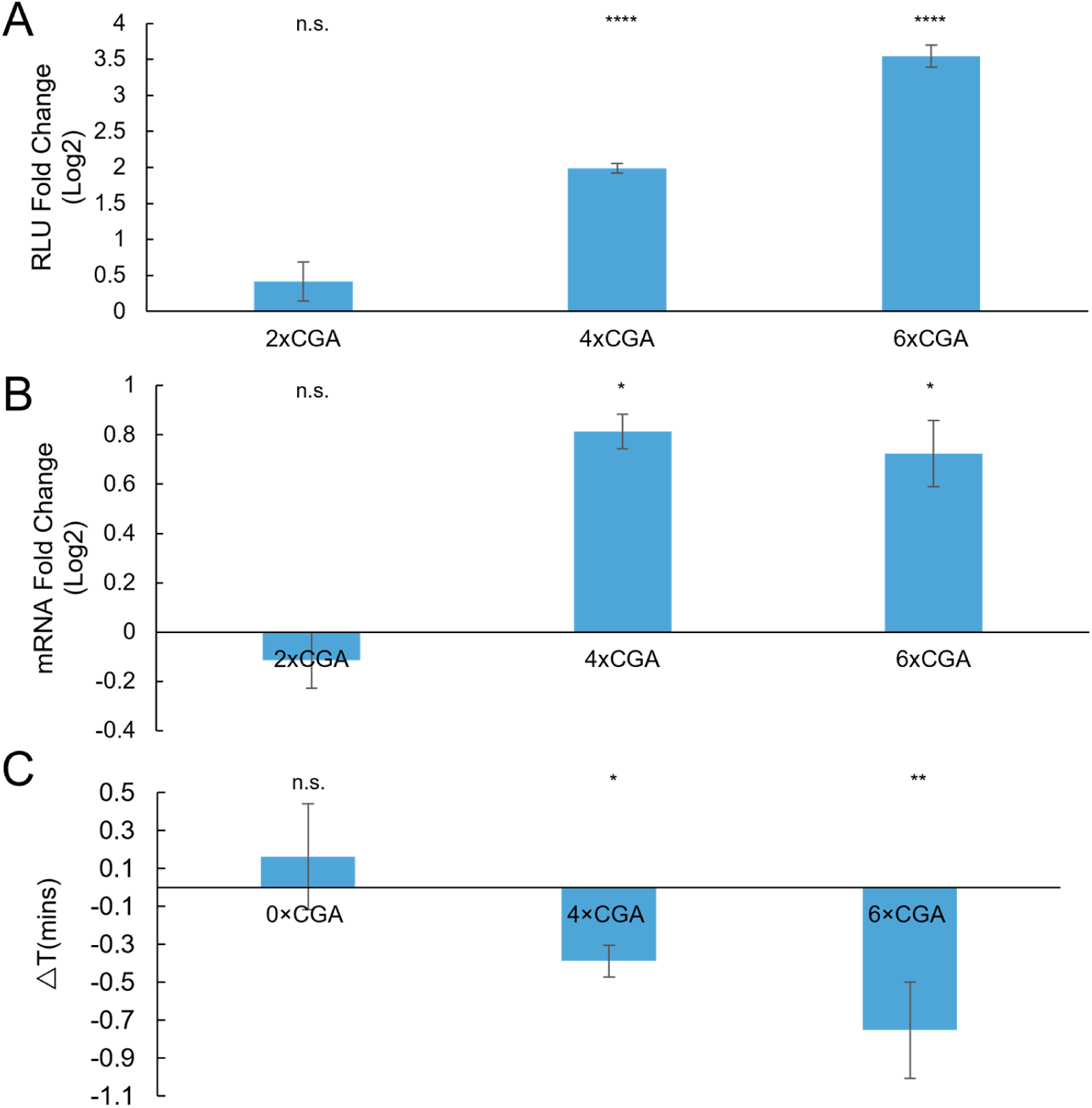
HEL2 deletion rescues protein expression, mRNA levels, and elongation time. A: Protein expression fold change of CGA constructs in a *hel2*∆ vs WT background (2xCGA n=2, 4xCGA n=7, 6xCGA n=7). B: mRNA level fold change of CGA constructs in a *hel2*∆ vs WT background (n = 3). C: Elongation delay of CGA constructs in a hel2∆ vs WT background (n = 3). All error bars indicate SEM. All statistical significances were calculated for each construct using two-tailed paired Student’s t-Test against

### Synonymous substitution to nonoptimal codons negatively impacts gene expression

Next, we asked how distributed slowdowns of non-optimal codons impact gene expression and elongation time. To study the impact of distributed non-optimal codons, we used our optYFP-nLuc construct and synonymously substituted the first 20 of 21 leucines for a nonoptimal leucine variant (Figure 4A, Supplementary Figure 3). First, we wanted to determine the impact of these synonymous substitutions on over-all elongation time. We measured the elongation time in each of our strains and compared it to the optimized strain to determine the elongation time delay associated with each synonymous substitution (Figure 4A). We found that substitution of the optimal leucine codon TTG for the nonoptimal codons CTC and CTT resulted in a significant delay in elongation time of approximately 0.5 and 2.5 minutes, respectively (Figure 4B). Due to the statistically significant differences in elongation time, we selected both the CTC and CTT-containing constructs for further study. Next, we measured the impact of codon substitution on protein and RNA levels (Figures 4C and 4D). As compared to the optimized control, we determined that substitution to the CTC codon reduced both protein and mRNA levels by approximately 20% and substitution to the CTT codon reduced both protein and mRNA levels by 50%. This was distinct from the RQC inducing CGA stalls that decreased protein production more substantially than they did mRNA levels.

**Fig. 4.**
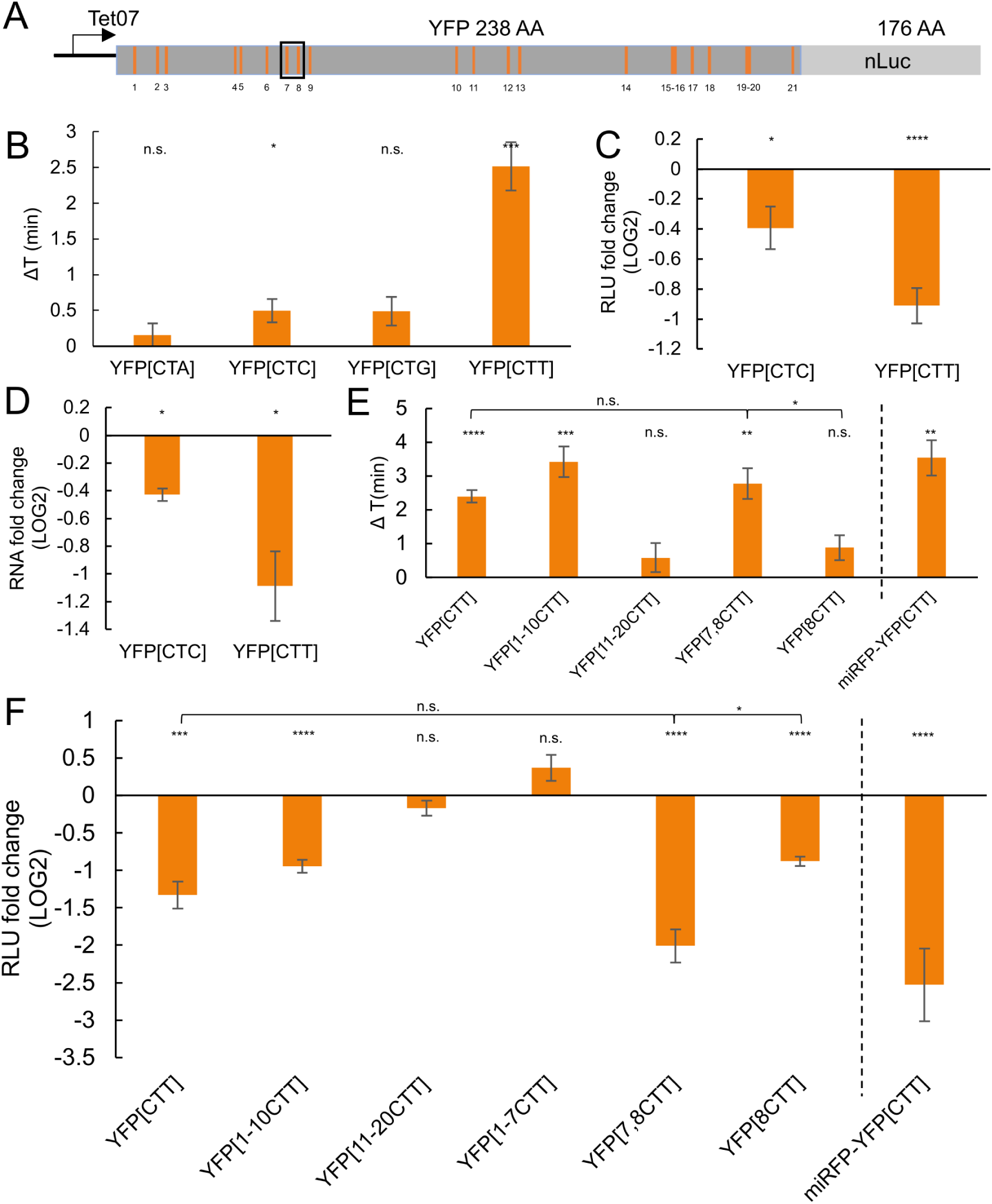
Distributed stalls in the YFP ORF decrease protein expression, mRNA levels, and delays elongation time. A: Schematic of position of 21 leucine codons in YFP-nLuc construct, with numbers from 5’ end to 3’ end labeled below. The black box emphasizes the positions of No.7 and No.8 Leu codons B: Elongation delay of distributed stall constructs compared to optYFP (YFP[CTA] n=6, YFP[CTC] n=6, YFP[CTG] n=5, YFP[CTT] n=7). The first 20 out of 21 total optimal TTG leucine codons in optYFP are synonymously substituted to a nonoptimal codon specified in brackets. C: Protein expression of distributed stall constructs normalized to optYFP control (n=4 for all). D: mRNA levels of distributed stall constructs normalized to optYFP control (n=3 for all). E: (Left) Elongation delay measurements of chimeric constructs normalized to optYFP control. (n=4 for YFP[CTT], YFP[7,8CTT] and YFP[8CTT]; n=6 for YFP[1-7CTT]; n=9 for YFP[1-10CTT] and YFP[11-20CTT]). (Right) Elongation delay measurements of miRFP-YFP[CTT] normalized to miRFP-optYFP control (n=6). F: (Left) Protein expression chimeric constructs normalized to optYFP control. (n= 3 for YFP[CTT], YFP[7,8CTT] and YFP[8CTT]; n=6 for YFP[1-7CTT], n=8 for YFP[1-10 CTT] and YFP[11-20CTT]). (Right) Protein expression of miRFP-YFP[CTT] normalized to miRFP-optYFP control.(n=6). All error bars indicate SEM. All statistical significances were calculated for each construct using two-tailed paired Student’s t-Test against optYFP control unless otherwise specified.

We sought to determine whether the increase in elongation time and decrease in protein expression observed was either contributed equally by each non-optimal codon or the specific placement of non-optimal codons in the YFP ORF. To assess this, we created a set of chimeric reporters in which the first 10 leucines in the YFP ORF were either optimal or nonoptimal followed by the next 10 leucines of the opposite optimality (Figure 4A). We hypothesized that if each codon contributed equally to elongation time, the elongation time delay of our chimeric constructs would be half of the delay between optYFP and YFP[CTT]. Instead, we found that both the elongation delay and protein expression of our chimeric YFP[1-10CTT] closely resembled YFP[CTT] and that our chimeric YFP[11-20CTT] closely resembled YFP[TTG] (Figures 4E and 4F). This provides evidence that substitution of leucines to a nonoptimal variant in the 5’ half of the YFP ORF is sufficient to drive protein expression and elongation time outcomes.

A previous study by Chu and colleagues showed that poor codons in the 5’ region of a transcript could negatively affect translation initiation through ribosome buildup preventing initiation from occurring, thereby reducing over-all translational output (34, 53). To test if the observed decrease in protein expression was a result of interference with initiation, we inserted a yeast-optimized miRFP (315 amino acids) upstream of our optYFP-nLuc and YFP[CTT]-nLuc constructs. We hypothesized that if initiation was negatively impacted by ribosome buildup, addition of a long yeast-optimized ORF upstream of the nonoptimal YFP[CTT] would rescue protein expression as compared to the optimal construct. Instead, we found that a statistically significant difference remained between the optimal and CTT-containing nonoptimal constructs (Figure 4F). Furthermore, we assessed the impact on elongation time and found that elongation time was not rescued to WT levels and the magnitude of delay is similar to the YFP[CTT] construct (Figure 4E). This suggests that the decrease in protein expression in the reporter is a result of the specific placement of the nonoptimal CTT codons within the 5’ half of the YFP ORF, but does not depend on the nonoptimal codons to be near the initiation codon.

To further dissect which leucine codons were important for the repression of protein expression, we made constructs that contained diminishing numbers of CTT codons from the first 10 leucine codons. We found that the first 8 and 9 codons showed similar levels of protein reduction but the first 7 leucine codons as CTT did not diminish expression (Figure 4F, Supplementary Figure 4B). As the switch took place from the 7th to 8th leucine codons, we tested whether the 8th leucine codon as a non-optimal CTT codon was sufficient to see the full effects. We found that while a single CTT at the 8th leucine codon significantly increased elongation time and decreased protein expression it was not to the same magnitude as the full YFP(CTT) construct (Figure 4E,F). We did however find that only two leucine codons, 7 and 8, switched to CTT were sufficient to fully repress protein expression and elongation time (Figure 4E,F). The switching of leucine 7 and 8 from TTG to CTT were associated with a more strongly folded local stem-loop structure as predicted by mFold (Supplementary Figure 4C).

Finally, we investigated if Hel2 or other translation sensors were responsible for the negative impacts on gene expression in our nonoptimal codon substituted constructs. Of particular interest was the RNA binding protein Dhh1, a conserved DEAD-box helicase previously shown to have roles in mRNA decapping and translational repression (54–57). Importantly, it has been shown to bind preferentially to mRNA with low codon optimality and has been proposed to slow down ribosome movement (32, 58). We hypothesized that the negative impacts on gene expression observed in YFP[CTC] and YFP[CTT] compared to the optYFP control may be a result of either Hel2 or Dhh1 influence. To test this, we transformed our optYFP, YFP[CTC], and YFP[CTT] constructs into either a *dhh1*Δ or *hel22*Δ strain.

First, we assessed the impact of protein expression on our constructs in a *dhh1*Δ or *hel2*Δ strain deletion background (Figure 5A). Based on Dhh1’s role in mediating translation repression of transcripts enriched in nonoptimal codons, we expected to see no impact in optYFP and a rescue of protein expression in YFP[CTC] and YFP[CTT]. Instead, we found differing effects for each construct; deletion of *DHH1* slightly increased protein expression in our optYFP construct, decreased protein expression in our YFP[CTC] construct, and had no statistically significant impact in our YFP[CTT] construct. We also found that *HEL2* deletion had no statistically significant effect on protein expression in any of our constructs (Figure 5A). This suggests that the drop in protein expression seen in the nonoptimal constructs was not due to a Hel2-mediated mechanism and is distinct from our acute CGA-containing constructs. Next, we examined the effect of Dhh1 on mRNA levels by comparing WT and *dhh1*Δ mRNA levels (Figure 5B). We found that deletion of *DHH1* decreased mRNA levels in our YFP[CTC] construct but had no statistically significant difference in the other constructs. The negative impact of *DHH1* deletion in our YFP[CTC] construct was of similar magnitude in protein and mRNA. This suggests that Dhh1 increases mRNA levels in our YFP[CTC] construct, which leads to increased protein expression.

**Fig. 5.**
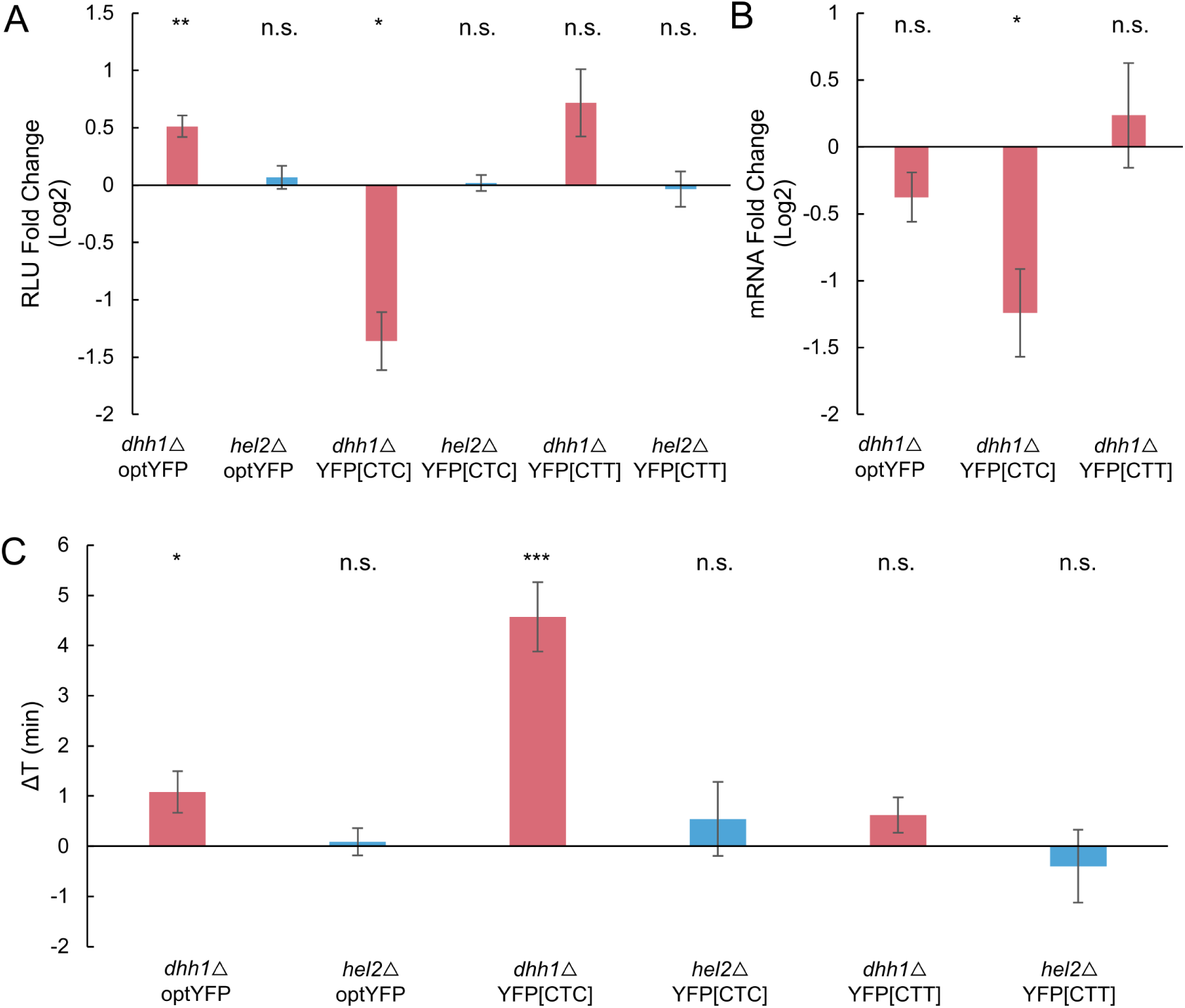
*DHH1* deletion, but not *HEL2* deletion, affects gene expression in substitution constructs. A: Protein expression of distributed stall constructs in *dhh1*∆ or *hel2*∆ background vs WT (n=5, 4, 5, 6, 7, and 3 from left to right). B: mRNA level fold change of distributed stall constructs in a *dhh1*∆ vs WT background (n = 5, 5, and 7 from left to right). C: Elongation delay of distributed stall constructs in *dhh1*∆ or *hel2*∆ background vs WT (n= 19, 4, 18, 6, 9, and 5 from left to right). All error bars indicate SEM. All statistical significances were calculated for each construct using two-tailed paired Student’s t-Test against WT control.

Lastly, we wanted to determine the impact of *dhh1*Δ or *hel2*Δ backgrounds on elongation time in our substituted leucine constructs. We measured elongation delay by comparing the elongation times of our constructs in each deletion strain to WT (Figure 5C). We found that deletion of *DHH1* slightly increased the elongation delay in our optYFP and more dramatically increased the elongation delay in the YFP[CTC] strain, suggesting that Dhh1 functions to speed up elongation in these constructs. However, we found no statistically significant difference in elongation time in our YFP[CTT] construct. Additionally, we found no statistically significant difference in elongation times in our *dhh1*Δ strains (Figure 5C). This is consistent with the *hel2*Δ strain protein expression data and supports the idea that a non-Hel2-mediated pathway is responsible for the negative impact on gene expression in our substituted leucine constructs.

## Discussion

Ribosome stalling and the connected quality control pathways are important for recognizing faulty and damaged mRNAs, yet quantitative measurements of how these stalls impact translation duration have been lacking. In this study, we developed a reporter assay to quantify the *in vivo* elongation time of various constructs containing stalling sequences in S. cerevisiae. Using CGA stalling reporters we find that total elongation time increases in a dose-dependent manner corresponding with the number of tandem CGA repeats while protein expression decreases logarithmically with increasing CGA repeats. Strikingly, we find that mRNA levels stabilize upon reaching a specific stall length, suggesting that the stall-activated NGD pathway reaches a maximum decay rate at 3x CGA. Interestingly the 50% reduction in mRNA levels is very similar to the mRNA reduction seen from a completely independently designed reporter containing 12xCGA (50) and other reporters containing 10xAAG (rare poly-Lysine) or 8xCCG (rare poly-proline codon) stalling sequences (22), further supporting NGD may be saturated at relatively shorter translational stalls.

From our synonymous leucine substitution constructs, we find that the nonoptimal codon CTT causes substantial delays in elongation time on the order of minutes, lengthening the elongation time of YFP approximately 3.5-fold. The elongation delay of 150s for the CTT reporter is similar to the elongation delay for our 4xCGA stalling reporter. Yet these two reporters behave very differently as the decrease in protein expression due to CTT could be explained completely by decreased mRNA levels, while the 4xCGA decreased protein levels to an even larger extent then the 50% decrease in mRNA levels. This pointed to the induction of RQC, which reduces protein expression on the CGA stalls through ribosome rescue. Further supporting this induction of RQC on CGA stalls but not CTT stalls, deletion of the RQC factor Hel2 could partially rescue the mRNA levels and protein production of CGA stalls, yet it had no significant effect on protein production and elongation times due to non-optimal CTT codons. These data point to further differentiation of ribosome stalling beyond just stall duration timing.

While initially 20 synonymous Leu codons were changed to poor CTT codons, not all non-optimal codons contribute equally to the elongation slowdown. Instead, we determined that two of the 20 codons were sufficient to drive the full elongation slowdown and repressed protein expression. This appears to be caused by local sequence effects and not specifically the poor codons being in the 5’ end of the ORF, as adding an upstream miRFP ORF was not able to rescue the translation slowdown and reduced protein production. This argues that local sequence context is important for determining the effects of codon optimality on gene expression. This fits with reports showing that specific combinations of codons modulate translation efficiency and mRNA decay (59, 60). We find that these two CTT substitutions drive the formation of a predicted stem-loop structure. Interestingly it was recently shown in prokaryotes that mRNA stem-loops can dock into the A site of the ribosome inhibiting translation elongation (61). It will be interesting to explore if a similar mechanism of translation elongation slowing takes place in this context.

We found that CGA stalls added 76s per CGA codon to the translation duration of the reporter after 3xCGAs. This led to an almost 5 minute lengthening of translation duration for a 6xCGA construct. A recent paper by Goldman and colleagues examined ribosomal clearance times on mRNA containing difficult-to-translate polyA-containing stretches and found it took approximately 10 and 13 minutes for ribosomes to clear off 50% of transcripts containing poly(A)36 and poly(A)60 stretches, respectively (27). Their finding on delays lasting on the order of minutes is consistent with our findings and represents an intriguing observation considering that the average half-life of yeast mRNAs is 10 minutes, suggesting that a significant portion of an mRNA’s half-life can be spent engaged in a ribosomal stall (62). The long duration of stalling also fits with long queues of ribosomes 5’ of the stall, as has been seen with disome-seq and in vivo translational imaging in mammalian cells (16, 27). We believe we may be observing a cumulative effect of ribosome queuing affecting the overall translation duration.

It is well-confirmed that Hel2 is a necessary factor mediating RQC and NGD pathways however its effects on ribosome stalling have been unclear. Two non-mutually exclusive models have been proposed (26, 52): first, since Hel2 is needed to promote the rescue of the stalled ribosome in a collision complex, deletion of *HEL2* will slow ribosome rescue, resulting in accumulated collided ribosomes, which increases elongation delay; in the second model, we propose that Hel2 is able to sense and stabilize stalled ribosomes to prevent further translation. In this scenario, deletion of *HEL2* would destabilize collided ribosomes, resulting in rescued elongation and shorter elongation delay. In this paper, we quantitatively measure the change of elongation delay after *HEL2* depletion and find a reduction in the translation duration of CGA stalled sequences. This is distinct from mammalian cells, where depletion of the mammalian homolog of Hel2, ZNF598, causes further delays in the clearing of ribosomes (27).

It has been previously reported that Dhh1 plays a role in degradation of mRNA enriched in nonoptimal codons (32). We were surprised to find that *DHH1* deletion instead decreases the expression of the YFP[CTC] construct. As the YFP constructs used in this study are all yeast optimized except for the leucine codons, it is possible that *DHH1* deletion would only be beneficial for mRNAs more enriched in poor codons. Previous work demonstrates a negligible effect of *dhh1*Δ on mRNA half-life for primarily optimal mRNA (32).

Although most studies have investigated Dhh1 with regards to its role in mRNA decay and translational repression, Dhh1 has also been shown to promote the translation of certain mRNAs. It has been previously demonstrated that a subset of mRNAs that contain highly-structured 5’UTRs and coding sequences require Dhh1 helicase activity for efficient expression (63) Furthermore, Dhh1 can shift roles in a condition-dependent manner. During nitrogen starvation, Dhh1 is required for the efficient expression of autophagy-related proteins Atg1 and Atg13, but when nutrients are plentiful Dhh1 encourages ATG mRNA degradation (64). Overall, this argues that Dhh1 may play context specific roles in translation elongation and may be able to speed up elongation in specific sequence contexts.

## Materials and methods

### Plasmid Preparation and Integration

All plasmids used in this study are listed in Supplementary Table S1. Plasmids containing synonymous leucine codon substituted YFP (TTG, CTA, CTC, CTG, and CTT) and a single-copy yeast integrating plasmid containing a *TET07* promoter were provided as a kind gift from Dr. Arvind Rasi Subramaniam at the Fred Hutchinson Cancer Center in Seattle, Washington. Fragments containing pTET07, YFP variants, and yeast-optimized nanoluciferase (Promega Cat. No. N1141) were amplified using PCR and cloned into the *XhoI* and *HindIII*-digested single-copy yeast integrating plasmid using Gibson assembly.

The pAG306-pTet07-YFP[1-7CTT]-nLuc, pAG306-pTet07-YFP[1-8CTT]-nLuc, pAG306-pTet07-YFP[1-9CTT]-nLuc, pAG306-pTet07-YFP[1-10CTT]-nLuc, pAG306-pTet07-YFP[11-20CTT]-nLuc strains were generated by PCR amplification of the entire backbone of the previous pAG306-pTet07-YFP[TTG]-nLuc plasmid beginning at nLuc and ending with pTet07, and PCR amplification of the corresponding parts of the YFP[TTG] and YFP [CTT] variants (For example, YFP[1-7CTT] includes No.1-7 Leu condon part of YFP[CTT] and No.8-20 Leu codon part of YFP[TTG]) (Supplementary Figure 4A). These fragments were combined using Gibson assembly.

The pAG306-pTet07-YFP[7,8CTT]-nLuc and pAG306-pTet07-YFP[8CTT] were further constructed with the same backbone of pAG306-pTet07-YFP[TTG]-nLuc plasmid, and distinct YFP variants. YFP[7,8CTT] and YFP[8CTT] variants are constructed based on YFP[TTG] and YFP[1-8CTT]. (For example, YFP[7,8CTT] includes No.1-6 Leu condon part of YFP[TTG] and No.7-20 Leu codon part of YFP[1-8CTT]) (Supplementary Figure 4A). These fragments were combined using Gibson assembly.

Plasmid variants containing two to six CGA stalls were generated using the aforementioned backbone PCR of the pAG306-pTet07-YFP[TTG] plasmid and a PCR amplified YFP[TTG] fragment containing two to six CGA repeats as a 3’ overhang. These fragments were combined using Gibson assembly.

All plasmids were linearized using *NotI* and integrated into yeast by homologous recombination. Integrations were screened by growing transformed yeast on synthetic complete (SC) dropout plates lacking uracil. These were then frozen down for long-term storage in YPD containing 15% v/v glycerol.

### Yeast Strains, Growth, and Media

The background yeast strain w303 (EY0690) was used for all experiments. Yeast *dhh1*Δ, *hel2*Δ, and *syh1*Δ strains were created by deleting the endogenous *DHH1, HEL2*, and *SYH1* loci, respectively, using pRS315 (Addgene Plasmid #3974) and screened by growing transformed yeast on SC dropout plates lacking leucine. Specific oligos used are listed in Supplementary Table S1. Yeast strains were frozen down in YPD containing 15% v/v glycerol.

For cells cultured for use in our reporter assay, cells were streaked out from frozen stocks onto YPD Agar plates and grown at 30 °C for two days. These plates were stored at 4 °C for up to one month.

### Luciferase-Based Elongation Reporter Assay

Liquid cultures were started from single colonies and allowed to grow overnight at 30 °C with shaking until an approximate OD600 of 0.3-0.5 after which cultures were divided into two tubes. For one of the tubes, 1 µL of a stock solution of anhydrotetracycline (250 µg/mL of ATC dissolved in EtOH) was added per mL of culture. Both tubes were returned to 30 °C with shaking for five to ten minutes. 90 µL of each culture was added to a 96-well white flat-bottom plate (Grainger) and to each well, 10 uL of furimazine (10 mM furimazine stock solution dissolved in DMSO diluted 1:200 in YPD), was added. Immediately after sample loading, the plate was placed in a 30 °C prewarmed Tecan Infinite® 200 PRO plate reader. The following program was used and luminescence measurements were taken every 30 or 60 seconds: (1) Kinetic Cycle: [Cycle Duration: 60 minutes, Kinetic Interval: 30 or 60 seconds], (2) Shaking: [Duration: 3 seconds, Mode: Orbital, Amplitude 2 mm], (3) Luminescence: [Attenuation: Automatic, Integration Time: 1000 ms, Settle Time: 0 ms].

### Schleif Plot and Elongation Delay Measurements

The Schleif Plot methodology was adapted from Schleif, R., et al. (1973) and slightly modified to assume a non-constant basal expression protein level. The general principle is that upon sufficient time for transcriptional induction to start there will be a proportional increase in mRNA levels to time (t). The increase of luciferase from a single mRNA is also proportional to time (t). As mRNA levels are also increasing with time this means that the total amount of luciferase is proportional to t2. For each sample, ATC-induced protein expression was calculated by subtracting the samples lacking ATC (-ATC) from the corresponding samples with ATC (+ATC) across all measured timepoints. Samples were then normalized to an OD600 of 1.0 by dividing their protein expression over time by their respective ODs. All values were then subtracted by the average RLU of the first 5 minutes to subtract background. Then, the square root of each value

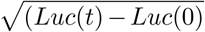

was calculated and plotted against time. Values that produced an error due to the square root of a negative value were set as “N/A” and avoided in our analysis. From this Schleif plot, we identified regions of linearity across our samples and selected a 10-15 minute window for analysis. Ideally, these regions of linearity are parallel between each sample and contain a minimal amount of noise. For each time window, we created a trendline and calculated the X-intercept of the trendline which represented the calculated elongation time of the sample. The calculated elongation time of the samples in a single assay were then compared to a control to determine elongation delay. These elongation delay measurements were then compared across assays and aggregated to determine the average elongation delay associated with the specific construct.

### RNA Extraction and Real Time qPCR

Yeast pellets were collected from samples 60-minutes post-ATC addition by spinning 1-1.5 mL of liquid culture at 3000 x g for 2 minutes and discarding the supernatant. These yeast pellets were then flash frozen in liquid nitrogen and stored at -80 °C until RNA extraction. RNA was extracted from yeast pellets using the MasterPure™ Yeast RNA Purification Kit (Lucigen Cat. No. MPY03100) according to the manufacturer’s instructions. RNA quality and concentration was assessed using a Nanodrop.

RNA samples were subjected to DNase digestion using RQ1 RNase-free DNase (Promega) according to the manufacturer’s instructions. cDNA was prepared from equal amounts of RNA from each sample using Protoscript II Reverse Transcriptase (New England Biolabs Cat. No. M0368X) and an oligo dT(18) primer according to the manufacturer’s instructions. RT-qPCR was done using a home-brew recipe with SYBR Green at a final concentration of 0.5X (Thermo Fisher S7564). Primers specific for nanoluciferase and actin are described in Supplementary Table S1. mRNA levels were normalized to ACT1 abundance and fold change was calculated by a standard Ct analysis.

## Acknowledgements

We would like to thank the Zid lab for helpful feedback on this manuscript. We thank Claes Andréasson for sharing the yeast optimized Nanoluciferase. This work was supported, in part, by the National Institutes of Health R35GM128798 (BMZ), funding from the UCSD Molecular Genetics Training grant (VH) and a training grant in Quantitative Integrative Biology from UCSD’s qBio Program (ATH)

