## Supplementary Figures for "Quantification of elongation stalls and impact on gene expression in yeast"

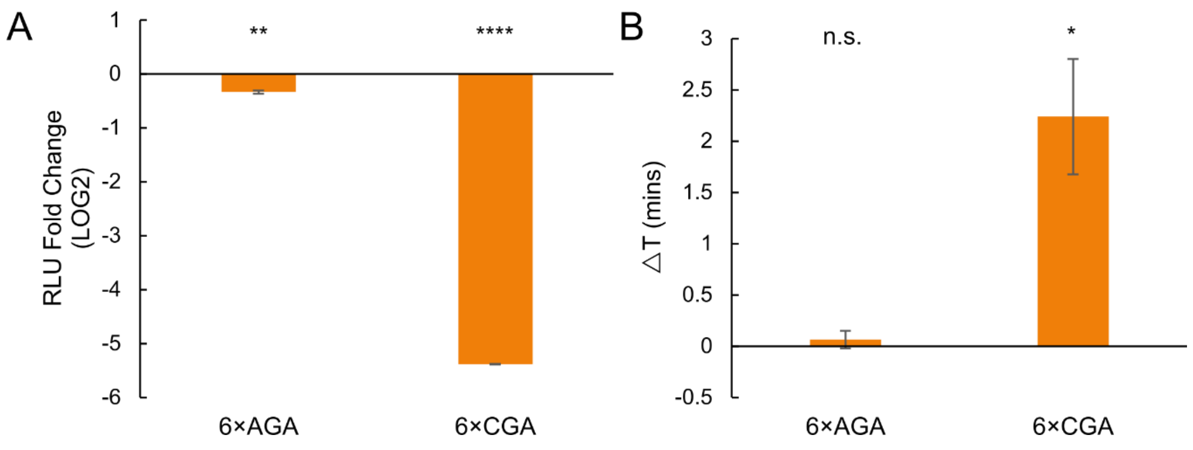


**Supplementary Figure 1. 6×CGA acute stall is induced by sequential non-optimal CGA codons, instead of its coding Arginine.**

A: Protein expression fold change of 6×AGA and 6×CGA constructs. (n=3 for both constructs)

B: Elongation change of 6×AGA and 6×CGA constructs vs. (n=3 for both constructs)

All error bars indicate SEM. All statistical significances were calculated for each construct using two-tailed paired Student's t-Test against optYFP.

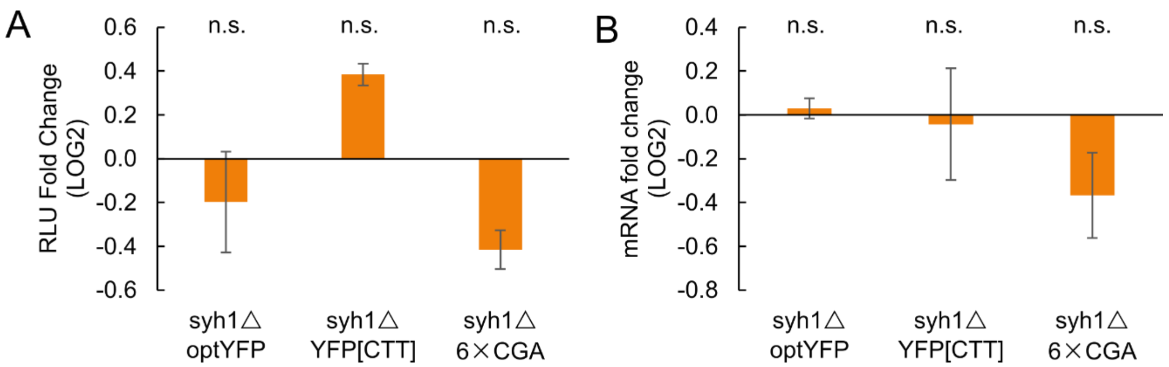


**Supplementary Figure 2. Syh1 doesn't significantly affects protein expression, mRNA expression and elongation of CTT and CGA-derived ribosome stalls**

A: Protein expression fold change of optYFP, YFP[CTT] and 6×CGA constructs in *syh1*△ vs WT (n=3 for all constructs)

B: mRNA expression fold change of optYFP, YFP[CTT] and 6×CGA constructs in *syh1*△ vs WT (n=3 for all constructs)

All error bars indicate SEM. All statistical significances were calculated for each construct using two-tailed paired Student's t-Test against WT constructs.

| Leu Codons | tAI | Construct |
|---|---|---|
| TTG (optimal) | 0.754 | optYFP |
| CTA | 0.185 | YFP[CTA] |
| CTC | 0.062 | YFP[CTC] |
| CTG | 0.059 | YFP[CTG] |
| CTT | 0.027 | YFP[CTT] |

**Supplementary Figure 3. Leucine codon variants.** Table of leucine codons used in this study. The optimal TTG codon is present in the optYFP control. Other constructs contain a nonoptimal leucine variant which is denoted in brackets. The tRNA Adaptation Index (tAI) is a metric that represents codon optimality and ranges from 0 (most nonoptimal) to 1 (most optimal).

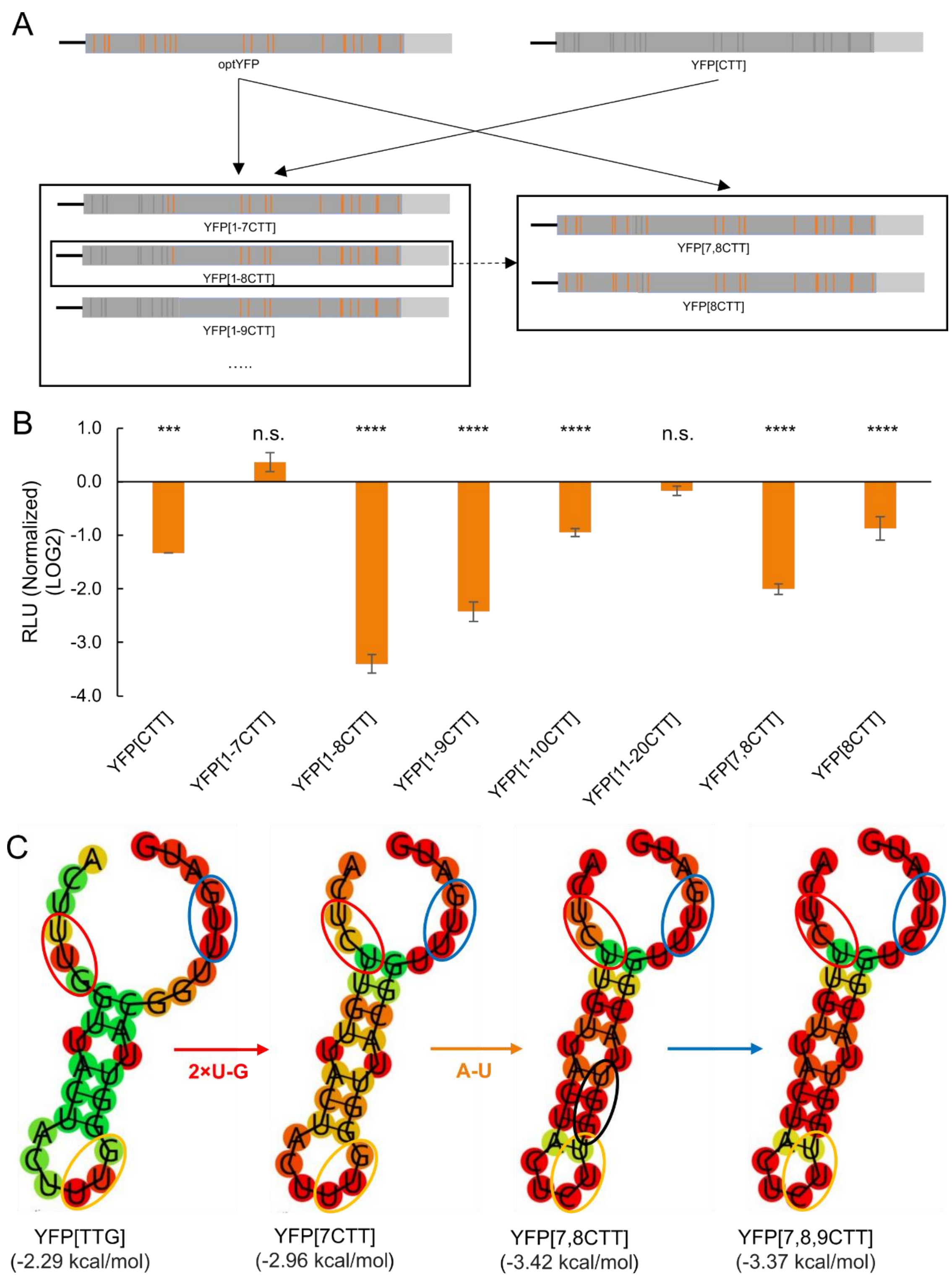


**Supplementary Figure 4.1 Cloning strategy of chimeric leucine constructs and mRNA structure induced distributed codon stalls**

A: Diagram of Chimeric YFP construction. YFP[1-7CTT], YFP[1-8CTT], YFP[1-9CTT], YFP[1-10CTT], YFP[10-20CTT] are constructed based on optYFP and YFP[CTT]; YFP[7,8CTT] and YFP[8CTT] are constructed based on optYFP and YFP[1-8CTT]

B: Protein expression of all chimeric leucine constructs. (n= 3 for YFP[CTT], YFP[1-8CTT], YFP[1-9CTT], YFP[7,8CTT] and YFP[8CTT]; n=6 for YFP[1-7CTT], n=8 for YFP[1-10 CTT] and YFP[11-20CTT])

C: RNA-fold prediction, with region from No.178 to No.207 bp

All error bars indicate SEM. All statistical significances were calculated for each construct using two-tailed paired Student's t-Test against optYFP.

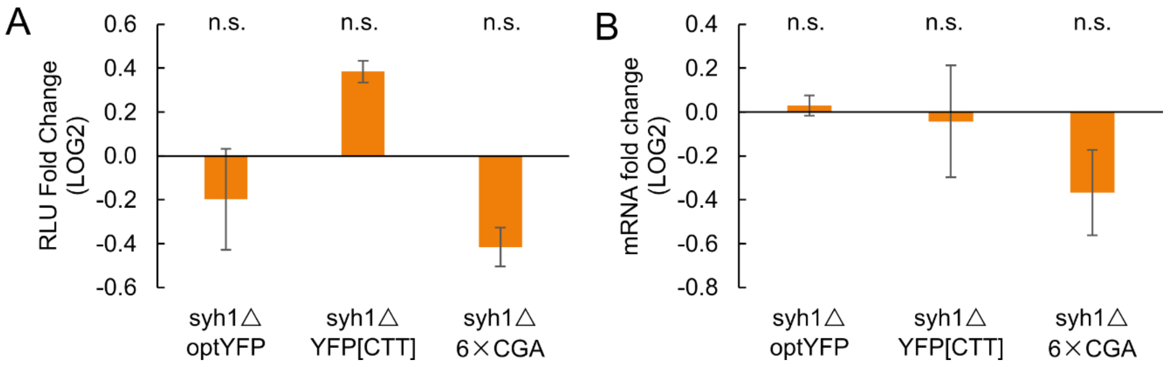


**Supplementary Figure 4 Syh1 doesn't significantly affects protein expression, mRNA expression and elongation of CTT and CGA-derived ribosome stalls**

A: Protein expression fold change of optYFP, YFP[CTT] and 6×CGA constructs in *syh1*△ vs WT (n=3 for all constructs)

B: mRNA expression fold change of optYFP, YFP[CTT] and 6×CGA constructs in *syh1*△ vs WT (n=3 for all constructs)

All error bars indicate SEM. All statistical significances were calculated for each construct using two-tailed paired Student's t-Test against WT constructs.
