## Supplementary Table S1 for "Quantification of elongation stalls and impact on gene expression in yeast"

| Identifier | Primer Name | Purpose | Sequence | PCR Temp |
|---|---|---|---|---|
| ZO1186 | Hel2 Deletion Fwd | Hel2 endogenous deletion | CTAATGCTATTGTCAGTTACAGGTTAGAAATATATTTCCAA CGG ATC CCC GGG TTA ATT AA | ZP191 |
| ZO1187 | Hel2 Deletion Rev | | CGAAAAAATAGTGGCTATACTTCTTTTCAAGAATTAGG GAA TTC GAG CTC GTT TAA AC | |
| ZO1192 | Hel2 Check Fwd | qPCR to check Hel2 endogenous deletion | TCTCAACCTACCTCAACTACC | |
| ZO1193 | Hel2 Check Rev | | GCTGCTTTTGTTTGCCTTTC | |
| ZO113 | Dhh1 Deletion Fwd | Dhh1 endogenous deletion | ATCCCAGGCCTAAAATACGACAAGAAAGAAAATAGTAGTA CGG ATC CCC GGG TTA ATT AA | ZP191 |
| ZO114 | Dhh1 Deletion Rev | | GCGTATCTCACCACAGTAGTTATTTTTTCTTAGATATTCT GAA TTC GAG CTC GTT TAA AC | |
| ZO131 | Dhh1 Check Fwd | qPCR to check Dhh1 endogenous deletion | ACAGCCGCATTTGTTATTCC | |
| ZO130 | Dhh1 Check Rev | | ACGACTTGGGAAGTTTGCAG | |
| ZO117 | Dom34 Deletion Fwd | Dom34 endogenous deletion | AAATGTAATTTAATGAAGATCCCAAAAAATTAAGCATTCG CGG ATC CCC GGG TTA ATT AA | ZP191 |
| ZO118 | Dom34 Deletion Rev | | AAATTTTATGTGTACATTACTTTTTTCTTACATAGTAAAT GAA TTC GAG CTC GTT TAA AC | |
| ZO127 | Dom34 Check Fwd | qPCR to check Dom34 endogenous deletion | AGAGCAATGGAGGAAAAGCA | |
| ZO126 | Dom34 Check Rev | | CCTCACCATCGTCTTCATCA | |
| ZO1331 | Syh1 Deletion Fwd | Syh1 endogenous deletion | TTTGCCACAGCTTGCACAAGATTGGCAGTGGCAGTAAGTG CGG ATC CCC GGG TTA ATT AA | ZP145 |
| ZO1332 | Syh1 Deletion Rev | | GCGTAGTAAACAACTACTATGGAACAAAAAGGCT GAA TTC GAG CTC GTT TAA AC | |
| ZO1335 | Syh1 Check Fwd | qPCR to check Syh1 endogenous deletion | TAATTTGGCGCCTTGGGCTA | |
| ZO1336 | Syh1 Check Rev | | AGATGGGGAAGGCGTTCTTG | |
| ZO83 | Actin Fwd | RT-qPCR | CTGCCGGTATTGACCAAACT | |
| ZO84 | Actin Rev | | CGGTGATTTCCTTTTGCATT | |
| ZO553 | nLuc Fwd | RT-qPCR | TGGTGATCAAATGGGTCAAA | |
| ZO544 | nLuc Rev | | CCTTCATAAGGACGACCAAA | |
| ZO1452 | pTet07_F | PCR pTet for ZP436 clone | GGAATTGACGAGTACGGTGGGTAGCTCGAG CCACTTCTAAATAAGCGAATTTC | ZP404 |
| ZO1453 | pTet07_R | | AATTGATCCGGTAATTTAGTGTG | |
| ZO1459 | yfp-nLucPEST_F | PCR nLuc for ZP436 clone | CACGGTATGGACGAATTGTACAAG ATGGTTTTTACTTTAGAAGATTTTG | ZP377 |
| ZO1463 | Cyc1term-nLuc_R | | GAATGTAAGCGTGACATAACTAATAAGCTTTTA AAAACCATGAGAATTAGCTAAAATACG | |
| ZO1455 | optYFP_F | PCR optYFP for ZP436 clone | ACTAAATTACCGGATCAATT ATGTCTAAGGGTGAAGAATTG | ZP408 |
| ZO1456 | optYFP_R | | TCTTCTAAAGTAAAAACCAT CTTGTACAATTCGTCCATAC | |
| ZO463 | NLuc+PestR | PCR pAG306 vector (including nLuc and pTet) | ATGGTTTTTACTTTAGAAGATTTTG | ZP436 |
| ZO1453 | pTet07_R | | AATTGATCCGGTAATTTAGTGTG | |
| ZO1464 | pTet-synYFP_F | PCR YFP for ZP432~ZP435 clone | ACTAAATTACCGGATCAATT ATGTCTAAGGGTGAAGAA | ZP405~ZP[illegible] |
| ZO1465 | nLuc-synYFP_R | | CAACAAAATCTTCTAAAGTAAAAACCAT CTTGTACAATTCGTCCATAC | |
| ZO1464 | pTet-synYFP_F | PCR YFP[TTG]-2CGA for ZP464 clone | ACTAAATTACCGGATCAATT ATGTCTAAGGGTGAAGAA | ZP408 |
| ZO1468 | 2xArgCGA-synYFP_R | | CAACAAAATCTTCTAAAGTAAAAACCAT TCGTCG CTTGTACAATTCGTCCATAC | |
| ZO1464 | pTet-synYFP_F | PCR YFP[TTG]-3CGA for ZP465 clone | ACTAAATTACCGGATCAATT ATGTCTAAGGGTGAAGAA | ZP408 |
| ZO1469 | 3xArgCGA-synYFP_R | | CAACAAAATCTTCTAAAGTAAAAACCAT TCGTCGTCG CTTGTACAATTCGTCCATAC | |
| ZO1464 | pTet-synYFP_F | PCR YFP[TTG]-4CGA for ZP466 clone | ACTAAATTACCGGATCAATT ATGTCTAAGGGTGAAGAA | ZP408 |
| ZO1470 | 4xArgCGA-synYFP_R | | CAACAAAATCTTCTAAAGTAAAAACCAT TCGTCGTCGTCG CTTGTACAATTCGTCCATAC | |

| | | | | |
|---|---|---|---|---|
| ZO1464 | pTet-synYFP_F | PCR YFP[TTG]-5CGA for ZP467 clone | ACTAAATTACCGGATCAATT ATGTCTAAGGGTGAAGAA | ZP408 |
| ZO1471 | 5xArgCGA-synYFP_R | | CAACAAAATCTTCTAAAGTAAAAACCAT TCGTCGTCGTCGTCG CTTGTACAATTCGTCCATAC | |
| ZO1464 | pTet-synYFP_F | PCR YFP[TTG]-6CGA for ZP468 clone | ACTAAATTACCGGATCAATT ATGTCTAAGGGTGAAGAA | ZP408 |
| ZO1472 | 6xArgCGA-synYFP_R | | CAACAAAATCTTCTAAAGTAAAAACCAT TCGTCGTCGTCGTCGTCG CTTGTACAATTCGTCCATAC | |
| ZO1464 | pTet-synYFP_F | PCR 5'part of YFP[1-7CTT] for ZP616 clone | ACTAAATTACCGGATCAATT ATGTCTAAGGGTGAAGAA | ZP434 |
| ZO1543 | LeuYFP-07_R | | CCGTAACCCAAAGTAGTAACAAGAGT | |
| ZO1542 | LeuYFP-07_F | PCR 3'part of YFP[1-7CTT] for ZP616 clone | ACTCTTGTTACTACTTTGGGTTACGG | ZP436 |
| ZO1465 | nLuc-synYFP_R | | CAACAAAATCTTCTAAAGTAAAAACCAT CTTGTACAATTCGTCCATAC | |
| ZO1464 | pTet-synYFP_F | PCR 5'part of YFP[1-8CTT] for ZP632 clone | ACTAAATTACCGGATCAATT ATGTCTAAGGGTGAAGAA | ZP434 |
| ZO1650 | LeuYFP-08_R V2 | | GCGAAACACATCAAACCGTAACC | |
| ZO1544 | LeuYFP-08_F | PCR 3'part of YFP[1-8CTT] for ZP632 clone | ACTCTTGTTACTACTCTTGGTTACGG | ZP436 |
| ZO1465 | nLuc-synYFP_R | | CAACAAAATCTTCTAAAGTAAAAACCAT CTTGTACAATTCGTCCATAC | |
| ZO1464 | pTet-synYFP_F | PCR 5'part of YFP[1-9CTT] for ZP617 clone | ACTAAATTACCGGATCAATT ATGTCTAAGGGTGAAGAA | ZP434 |
| ZO1560 | LeuYFP-09_R | | GGCATAGCAGACTTGAAGAAGTCG | |
| ZO1559 | LeuYFP-09_F | PCR 3'part of YFP[1-9CTT] for ZP617 clone | CGACTTCTTCAAGTCTGCTATGCC | ZP436 |
| ZO1465 | nLuc-synYFP_R | | CAACAAAATCTTCTAAAGTAAAAACCAT CTTGTACAATTCGTCCATAC | |
| ZO1464 | pTet-synYFP_F | PCR 5'part of YFP[1-10CTT] for ZP513 clone | ACTAAATTACCGGATCAATT ATGTCTAAGGGTGAAGAA | ZP434 |
| ZO1486 | FirstHalfYFP_R | | CCGTCTTCCTTGAAGTCGATACCC | |
| ZO1487 | SecondHalfYFP_F | PCR 3'part of YFP[1-10CTT] for ZP513 clone | GGGTATCGACTTCAAGGAAGACGG | ZP436 |
| ZO1465 | nLuc-synYFP_R | | CAACAAAATCTTCTAAAGTAAAAACCAT CTTGTACAATTCGTCCATAC | |
| ZO1464 | pTet-synYFP_F | PCR 5'part of YFP[11-20CTT] for ZP515 clone | ACTAAATTACCGGATCAATT ATGTCTAAGGGTGAAGAA | ZP436 |
| ZO1486 | FirstHalfYFP_R | | CCGTCTTCCTTGAAGTCGATACCC | |
| ZO1487 | SecondHalfYFP_F | PCR 3'part of YFP[11-20CTT] for ZP515 clone | GGGTATCGACTTCAAGGAAGACGG | ZP434 |
| ZO1465 | nLuc-synYFP_R | | CAACAAAATCTTCTAAAGTAAAAACCAT CTTGTACAATTCGTCCATAC | |
| ZO1464 | pTet-synYFP_F | PCR 5'part of YFP[7,8CTT] for ZP634 clone | ACTAAATTACCGGATCAATT ATGTCTAAGGGTGAAGAA | ZP436 |
| ZO1652 | LeuYFP[7,8 CTT]_R | | AGTTGGCCATGGAACTGG | |
| ZO1651 | LeuYFP[7,8 CTT]_F | PCR 3'part of YFP[7,8CTT] for ZP634 clone | CCAGTTCCATGGCCAACT | ZP632 |
| ZO1465 | nLuc-synYFP_R | | CAACAAAATCTTCTAAAGTAAAAACCAT CTTGTACAATTCGTCCATAC | |
| ZO1464 | pTet-synYFP_F | PCR 5'part of YFP[8CTT] for ZP635 clone | ACTAAATTACCGGATCAATT ATGTCTAAGGGTGAAGAA | ZP436 |
| ZO1654 | LeuYFP[8 CTT]_R | | ACCGTAACCAAGAGTAGTAAC | |
| ZO1653 | LeuYFP[8 CTT]_F | PCR 3'part of YFP[8CTT] for ZP635 clone | GTTACTACTCTTGGTTACGGT | ZP632 |
| ZO1465 | nLuc-synYFP_R | | CAACAAAATCTTCTAAAGTAAAAACCAT CTTGTACAATTCGTCCATAC | |
| ZO1464 | pTet-synYFP_F | PCR 5'part of YFP[7,8CTT](186T>A) for ZP644 c | ACTAAATTACCGGATCAATT ATGTCTAAGGGTGAAGAA | ZP634 |
| ZO1679 | YFP point mutation (ACT to | | CAAACCGTAACCAAGAGTTGTAAC | |
| ZO1678 | YFP point mutation (ACT to | PCR 3'part of YFP[7,8CTT](186T>A) for ZP644 c | GTTACAACTCTTGGTTACGGTTTG | ZP634 |
| ZO1465 | nLuc-synYFP_R | | CAACAAAATCTTCTAAAGTAAAAACCAT CTTGTACAATTCGTCCATAC | |
| ZO1464 | pTet-synYFP_F | PCR 5'part of YFP[7,8CTT](195T>C) for ZP645 c | ACTAAATTACCGGATCAATT ATGTCTAAGGGTGAAGAA | ZP634 |

| | | | | |
|---|---|---|---|---|
| ZO1681 | YFP point mutation (GGT to | PCR 5'part of YFP[7,8CTT](195T>C) for ZP645 c | CAAACCGTAGCCAAGAGTAGTAAC | ZP634 |
| ZO1680 | YFP point mutation (GGT to | PCR 3'part of YFP[7,8CTT](195T>C) for ZP645 c | GTTACTACTCTTGGCTACGGTTTG | ZP634 |
| ZO1465 | nLuc-synYFP_R | | CAACAAAATCTTCTAAAGTAAAAACCAT CTTGTACAATTCGTCCATAC | |
| ZO1464 | pTet-synYFP_F | PCR 5'part of YFP[7,8CTT](186T>A, 195T>C) fo | ACTAAATTACCGGATCAATT ATGTCTAAGGGTGAAGAA | ZP634 |
| ZO1683 | YFP point mutation (ACT to | | CAAACCGTAGCCAAGAGTTGTAAC | |
| ZO1682 | YFP point mutation (ACT to | PCR 3'part of YFP[7,8CTT](186T>A, 195T>C) fo | GTTACAACTCTTGGCTACGGTTTG | ZP634 |
| ZO1465 | nLuc-synYFP_R | | CAACAAAATCTTCTAAAGTAAAAACCAT CTTGTACAATTCGTCCATAC | |
| ZO1493 | pTet-miRFP_F | PCR miRFP for ZP530/ZP531 | ACTAAATTACCGGATCAATT ATGGTAGCAGGCCATGCAAG | ZP317 |
| ZO1494 | synYFP_miRFP_R | | TTCTTCACCCTTAGACAT GCTTTCCAGAGCTGTAATCC | |

**late**

| Designa | Strain name | Reagent type (species) | Source | Identifiers |
|---|---|---|---|---|
| ZY8 | ZY8 | Saccharomyces cerevisiae | https://www.yeastgenome.org/strain/S00020345 | BY4741 |
| ZY549 | ZY8 hel2△ | Saccharomyces cerevisiae | This study | BY4741, hel2△ |
| ZY501 | ZY8 dhh1 △ | Saccharomyces cerevisiae | This study | BY4741, dhh1△ |
| ZY502 | ZY8 dom34 △ | Saccharomyces cerevisiae | This study | BY4741, dom34△ |
| ZY803 | ZY8 syh1 △ | Saccharomyces cerevisiae | This study | BY4741, syh1△ |
| ZY478 | nLuc | ZY8 background | This study | BY4741, pAG306-ura-rtta-pTet-nLuc |
| ZY483 | optYFP | | This study | BY4741, pAG306-ura-rtta-pTet-YFP[TTG]-nLuc |
| ZY532 | 2xCGA | | This study | BY4741, pAG306-ura-rtta-pTet-YFP[TTG]-2xCGA-nLuc |
| ZY578 | 3xCGA | | This study | BY4741, pAG306-ura-rtta-pTet-YFP[TTG]-3xCGA-nLuc |
| ZY533 | 4xCGA | | This study | BY4741, pAG306-ura-rtta-pTet-YFP[TTG]-4xCGA-nLuc |
| ZY579 | 5xCGA | | This study | BY4741, pAG306-ura-rtta-pTet-YFP[TTG]-5xCGA-nLuc |
| ZY534 | 6xCGA | | This study | BY4741, pAG306-ura-rtta-pTet-YFP[TTG]-6xCGA-nLuc |
| ZY836 | 6xAGA | | This study | BY4741, pAG306-ura-rtta-pTet-YFP[TTG]-6xAGA-nLuc |
| ZY484 | YFP[CTA] | | This study | BY4741, pAG306-ura-rtta-pTet-YFP[CTA]-nLuc |
| ZY480 | YFP[CTC] | | This study | BY4741, pAG306-ura-rtta-pTet-YFP[CTC]-nLuc |
| ZY481 | YFP[CTT] | | This study | BY4741, pAG306-ura-rtta-pTet-YFP[CTT]-nLuc |
| ZY482 | YFP[CTG] | | This study | BY4741, pAG306-ura-rtta-pTet-YFP[CTG]-nLuc |
| ZY870 | YFP[1-7CTT] | | This study | BY4741, pAG306-ura-rtta-pTet-YFP[1-7CTT]-nLuc |
| ZY871 | YFP[1-8CTT] | | This study | BY4741, pAG306-ura-rtta-pTet-YFP[1-8CTT]-nLuc |
| ZY872 | YFP[1-9CTT] | | This study | BY4741, pAG306-ura-rtta-pTet-YFP[1-9CTT]-nLuc |
| ZY624 | YFP[1-10CTT] | | This study | BY4741, pAG306-ura-rtta-pTet-YFP[1-10CTT]-nLuc |
| ZY626 | YFP[11-20CTT] | | This study | BY4741, pAG306-ura-rtta-pTet-YFP[11-20CTT]-nLuc |
| ZY877 | YFP[7,8CTT] | | This study | BY4741, pAG306-ura-rtta-pTet-YFP[7,8CTT]-nLuc |
| ZY878 | YFP[8CTT] | | This study | BY4741, pAG306-ura-rtta-pTet-YFP[8CTT]-nLuc |
| ZY897 | 186T>A | | This study | BY4741, pAG306-ura-rtta-pTet-YFP[7,8CTT] (186T>A)-nLuc |
| ZY898 | 195T>C | | This study | BY4741, pAG306-ura-rtta-pTet-YFP[7,8CTT] (195T>C)-nLuc |
| ZY899 | 186T>A, 195T>C | | This study | BY4741, pAG306-ura-rtta-pTet-YFP[7,8CTT] (186T>A, 195T>C)-nLuc |
| ZY692 | miRFP-optYFP | | This study | BY4741, pAG306-ura-rtta-pTet-miRFP-YFP[TTG]-nLuc |
| ZY691 | miRFP-YFP[CTT] | | This study | BY4741, pAG306-ura-rtta-pTet-miRFP-YFP[CTT]-nLuc |
| ZY553 | optYFP hel2△ | Saccharomyces cerevisiae (ZY | This study | BY4741, hel2△, pAG306-ura-rtta-pTet-YFP[TTG]-nLuc |
| ZY566 | 2xCGA hel2△ | | This study | BY4741, hel2△, pAG306-ura-rtta-pTet-YFP[TTG]-2xCGA-nLuc |
| ZY567 | 4xCGA hel2△ | | This study | BY4741, hel2△, pAG306-ura-rtta-pTet-YFP[TTG]-4xCGA-nLuc |

| | | | | |
|---|---|---|---|---|
| ZY568 | 6xCGA hel2△ | Saccharomyces cerevisiae (ZY | This study | BY4741, hel2△, pAG306-ura-rtta-pTet-YFP[TTG]-6xCGA-nLuc |
| ZY550 | YFP[CTC] hel2△ | | This study | BY4741, hel2△, pAG306-ura-rtta-pTet-YFP[CTC]-nLuc |
| ZY551 | YFP[CTT] hel2△ | | This study | BY4741, hel2△, pAG306-ura-rtta-pTet-YFP[CTT]-nLuc |
| ZY519 | optYFP dhh1△ | Saccharomyces cerevisiae (ZY | This study | BY4741, dhh1△, pAG306-ura-rtta-pTet-YFP[TTG]-nLuc |
| ZY516 | YFP[CTC] dhh1△ | | This study | BY4741, dhh1△, pAG306-ura-rtta-pTet-YFP[CTC]-nLuc |
| ZY522 | YFP[CTT] dhh1△ | | This study | BY4741, dhh1△, pAG306-ura-rtta-pTet-YFP[CTT]-nLuc |
| ZY520 | optYFP dom34△ | Saccharomyces cerevisiae (ZY | This study | BY4741, dom34△, pAG306-ura-rtta-pTet-YFP[TTG]-nLuc |
| ZY517 | YFP[CTC] dom34△ | | This study | BY4741, dom34△, pAG306-ura-rtta-pTet-YFP[CTC]-nLuc |
| ZY523 | YFP[CTT] dom34△ | | This study | BY4741, dom34△, pAG306-ura-rtta-pTet-YFP[CTT]-nLuc |
| ZY808 | optYFP syh1△ | Saccharomyces cerevisiae (ZY | This study | BY4741, syh1△, pAG306-ura-rtta-pTet-YFP[TTG]-nLuc |
| ZY809 | 6xCGA syh1△ | | This study | BY4741, syh1△, pAG306-ura-rtta-pTet-YFP[TTG]-6xCGA-nLuc |
| ZY807 | YFP[CTT] syh1△ | | This study | BY4741, syh1△, pAG306-ura-rtta-pTet-YFP[CTT]-nLuc |

| Designat | Construct Name | Source | Identifiers | Additional information |
|---|---|---|---|---|
| ZP404 | pLLSC15 | Arvind (Rasi) Subramania | pAG306-ura-rtta-yfpCTA | Parent plasmid to PCR YFP[CTA] |
| ZP405 | pLLSC16 | Arvind (Rasi) Subramania | pAG306-ura-rtta-yfpCTC | Parent plasmid to PCR YFP[CTC] |
| ZP406 | pLLSC17 | Arvind (Rasi) Subramania | pAG306-ura-rtta-yfpCTT | Parent plasmid to PCR YFP[CTT] |
| ZP407 | pLLSC18 | Arvind (Rasi) Subramania | pAG306-ura-rtta-yfpCTG | Parent plasmid to PCR YFP[CTG] |
| ZP408 | pLLSC19 | Arvind (Rasi) Subramania | pAG306-ura-rtta-yfpwt[TTG/AGA] | Parent plasmid to PCR optYFP, source of pTet and pAG306 vector |
| ZP377 | nLuc source | Lab stock | TetO7-PGK1UTR-LacZ-PKTlinker-nLucPEST-MS2(v4) | Parent plasmid to PCR nLuc |
| ZP317 | miRFPsource | Lab stock | Hsp30pr-Hsp26UTR-nLuc-pKTlinker-miRFPPEST-12xMS2v6-ADH1ter | Parent plasmid to PCR miRFP |
| ZP191 | Pringle (HPH) | Wilhelm lab | pFA6a-HphMX6 | PCR HPH marker for endogenous deletion |
| ZP145 | Pringle (KAN) | Lab stock | pFA6a-Kan | PCR KAN marker for endogenous deletion |
| ZP427 | nLuc | This study | pAG306-ura-rtta-pTet-nLuc | |
| ZP436 | optYFP | This study | pAG306-ura-rtta-pTet-YFP[TTG]-nLuc | |
| ZP464 | 2xCGA | This study | pAG306-ura-rtta-pTet-YFP[TTG]-2xCGA-nLuc | |
| ZP486 | 3xCGA | This study | pAG306-ura-rtta-pTet-YFP[TTG]-3xCGA-nLuc | |
| ZP465 | 4xCGA | This study | pAG306-ura-rtta-pTet-YFP[TTG]-4xCGA-nLuc | |
| ZP487 | 5xCGA | This study | pAG306-ura-rtta-pTet-YFP[TTG]-5xCGA-nLuc | |
| ZP466 | 6xCGA | This study | pAG306-ura-rtta-pTet-YFP[TTG]-6xCGA-nLuc | |
| ZP599 | 6xAGA | This study | pAG306-ura-rtta-pTet-YFP[TTG]-6xAGA-nLuc | |
| ZP432 | YFP[CTA] | This study | pAG306-ura-rtta-pTet-YFP[CTA]-nLuc | |
| ZP433 | YFP[CTC] | This study | pAG306-ura-rtta-pTet-YFP[CTC]-nLuc | |
| ZP434 | YFP[CTT] | This study | pAG306-ura-rtta-pTet-YFP[CTT]-nLuc | |
| ZP435 | YFP[CTG] | This study | pAG306-ura-rtta-pTet-YFP[CTG]-nLuc | |
| ZP616 | YFP[1-7CTT] | This study | pAG306-ura-rtta-pTet-YFP[1-7CTT]-nLuc | |
| ZP632 | YFP[1-8CTT] | This study | pAG306-ura-rtta-pTet-YFP[1-8CTT]-nLuc | |
| ZP617 | YFP[1-9CTT] | This study | pAG306-ura-rtta-pTet-YFP[1-9CTT]-nLuc | |
| ZP513 | YFP[1-10CTT] | This study | pAG306-ura-rtta-pTet-YFP[1-10CTT]-nLuc | |
| ZP515 | YFP[11-20CTT] | This study | pAG306-ura-rtta-pTet-YFP[11-20CTT]-nLuc | |
| ZP634 | YFP[7,8CTT] | This study | pAG306-ura-rtta-pTet-YFP[7,8CTT]-nLuc | |
| ZP635 | YFP[8CTT] | This study | pAG306-ura-rtta-pTet-YFP[8CTT]-nLuc | |
| ZP644 | 186T>A | This study | pAG306-ura-rtta-pTet-YFP[7,8CTT] (186T>A)-nLuc | |
| ZP645 | 195T>C | This study | pAG306-ura-rtta-pTet-YFP[7,8CTT] (195T>C)-nLuc | |
| ZP646 | 186T>A, 195T>C | This study | pAG306-ura-rtta-pTet-YFP[7,8CTT] (186T>A, 195T>C)-nLuc | |
| ZP531 | miRFP-optYFP | This study | pAG306-ura-rtta-pTet-miRFP-YFP[TTG]-nLuc | |
| ZP530 | miRFP-YFP[CTT] | This study | pAG306-ura-rtta-pTet-miRFP-YFP[CTT]-nLuc | |
